# To examine environmental pollution by economic growth and their impact in an environmental Kuznets curve (EKC) among developed and developing countries

**DOI:** 10.1101/492348

**Authors:** YuSheng Kong, Rabnawaz Khan

## Abstract

This study analyzes the core energy consumption among countries specific variables by Environmental Kuznets Curve hypothesis (EKC), for a panel data of 29 (14 developed and 15 developing) countries during the period of 1977-2014. By assessing Generalized Method of Moments (GMM) regressions with first generation test such as common root, individual Augmented Dickey-Fuller (ADF), and individual root-Fisher-PP have been computed individually, the results confirm the EKC hypothesis in the case of emissions of solid, liquid, gases, manufacturing industries and also construction. Hence, we computed the cointegration test by Pedroni Kao from Engle-Granger based and Fisher. Onward, since the variable are co-integrated, a panel vector error correction model is estimated in GDP per capita, emission from manufacturing industries, arms import, commercial service export and coal rent, order to perform Pairwise Granger Causality test and indicate Vector Error Correction (VEC), with co-integration restrictions. Moreover, the statistical finding from VEC short-run unidirectional causality from GDP per capita growth to manufacturing industries and coat rent, as well as the causal link with manufacturing industries and commercial service export. Additionally, since there occurred no causal link among economic growth, arm import and coal rent.

## 1. Introduction

Developing countries, with rapid development of economy are leading the growth of energy consumption globally.^1^ The energy consumption of developing countries was 7.64×10^9^ (ton) oil equivalent (toe), accounting for 58.1% in 2005 all over the world, also in 2015 the consumption of energy increase in developing countries by 2.38×10^9^ (toe)^2^. The level of energy intensity in China (8.34), Russia (9.49) and Germany (3.88)^3^, indicate big gap between developing and developed countries. Other side the developing countries decrease energy intensity slowly and try to achieve the bottleneck problems with well-developed technology. Furthermore, 79% developed countries are responsible for historical carbon emission, in which USA is 22%, European Union is 40% and China is 9%^4^ Fig-1. There are 60% of CO2 emission responsible countries are China and USA, it’s is two-fifth and these top polluters do about the heat-trapping gases liable for global warming and their infections.^5^ Also in 2013 CO2 emission is 11 billion tons with 1.36 billion population. The 62% coal consumption cap has been announced by 2020 in China.^6^

**Fig-1:**
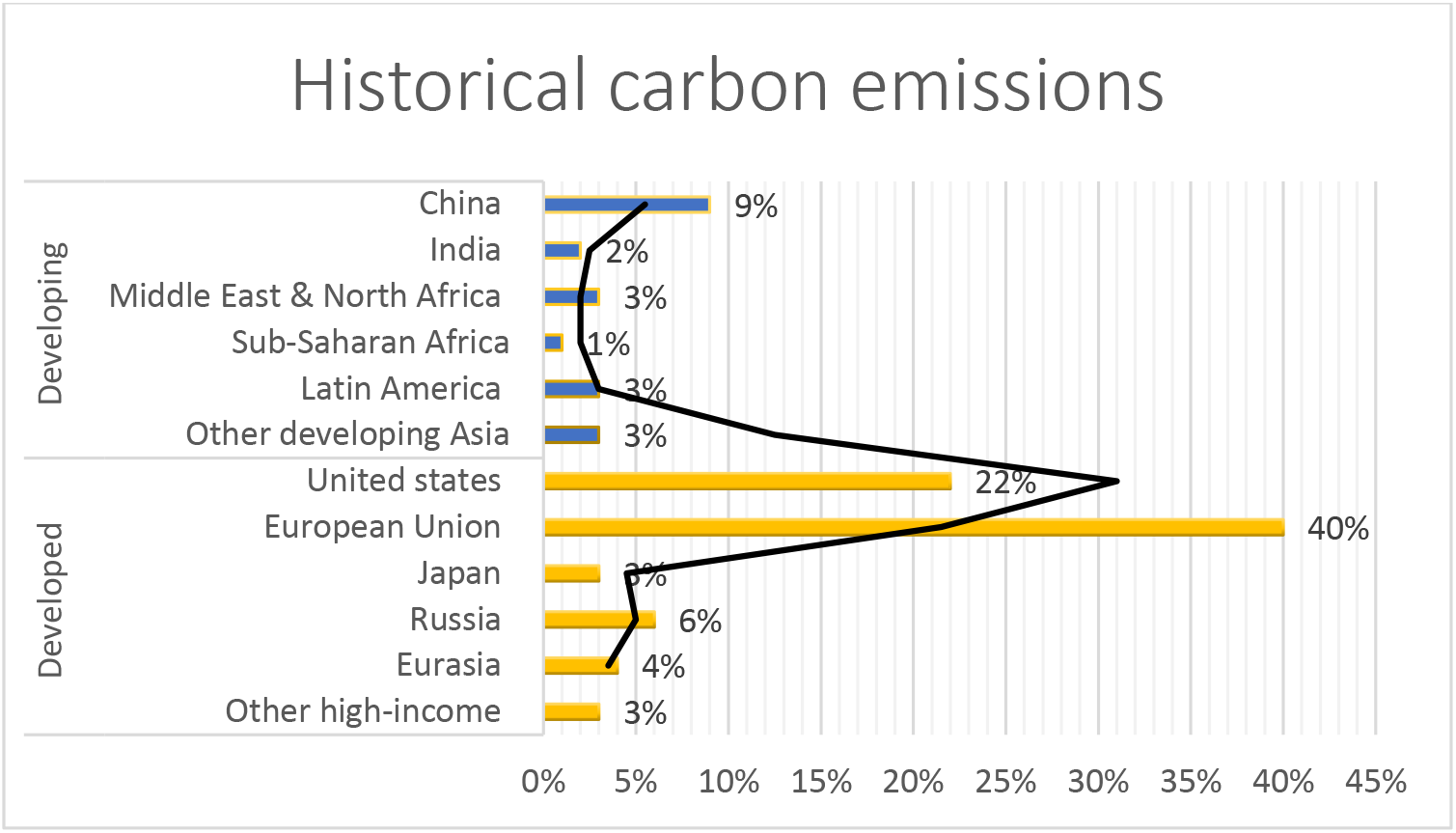
Historical carbon emission. Source: LUCEF, 1850-2011(CAITv2.0

The solid fuel consumption varies in different countries regarding with magnitude of indicators, the darker shade and higher the value. The China highest value in all over the world is 7,431,146.00. The Bolivia is the lowest value with 0.00.^7^ CO2 is naturally occurred with gas fixed by photosynthesis into organic matter, also biomass burning and byproduct of fuel consumption of fossil emitted from land use to changes along with industrial processes. The industrial revolution has rapidly increased global warming and atmospheric carbon dioxide. Burning wood, oil, coal and waste material, such as in industrial process of cement has been increased CO2 emission.^8^

The USA is one of top developed country by CO2 emission from gaseous fuel consumption in all over the world and 1.43 million kt that account for 21.72% of world’s CO2 emission from gaseous fuel consumption in 2014. Other five top countries (China, Russian Federation, Iran, and Japan), 48.97% account of it. In 2014, estimated emission of CO2 from fuel gaseous was at 6.6 million.^9^ Furthermore, it’s injected to the melting zone, auto-ignited (Solid combustion zone) and the methane concentrations of 0 to 5% vol, also the total calorific heat input unchanged. The pattern of heat in melting zone was recorded by non-contact thermal infrared imager and thermocouples. Significantly, the result indicated that extend the melting zone from the upstream and it higher than from coke sintering, without increasing the energy consumption. Therefore, the saving potential was evaluated by reducing the heat 4 to 8%. [1]. The continue modification and well-developed technology have been directly affected on solid combustion zone, like 15% energy consumption in iron and steel industry in the China and 26% consumption in pre-treatment process. The CH4 emission were approximately 5.1million tones, equivalent to 10.78 million of CO2, it indicated the third largest source of CH4 emission.

Municipal solid waste (MSW) landfills 69% of the solid waste which received from USA (94% of total landfills emission). Furthermore, the waste of energy emission was accounted 12.1 million metric tonnes of CO2 emission competitively 1745 million emitted in the field of transportation.

While 26.5 million tonnes incineration is used to treat of waste in USA, or approx. 7 to 19 percent solid waste generated. Meanwhile, 3.2% CO2 emission have been increased in 2010 and total greenhouse gases were equivalent to 6.82% billion metric tonnes of CO2^10^. While CO2 is found in our environment but the problem is that the industrial revolution has increased the quantity of it in 19^th^ century by industrial modification Fig-2, because it’s most prominent greenhouse gases climate change^11^ and most of the scientists agree on that is not only for Chinese hoax.^12^

**Fig-2:**
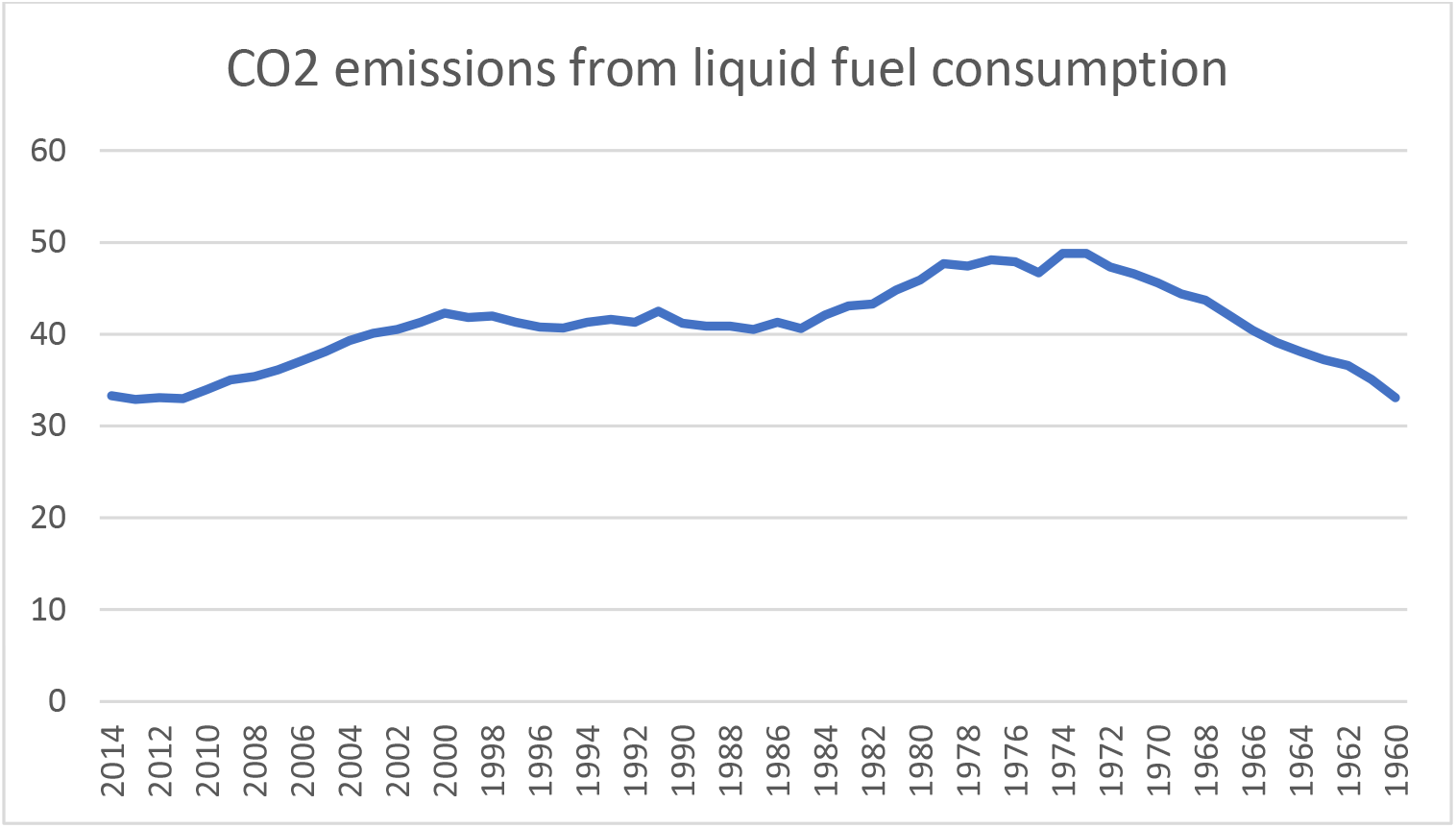
CO2 emission from liquid fuel consumption. Source: Authors ‘ amplification.

Environmental Kuznets Curve (EKC) have been already explored different ideas in CO2 emission but there are not too many studies for pollutant emission in developed and developing countries in period of 1977 to 2014. In this research EKC hypothesis for CO2 emission from gasses, liquid, solid fuel, and manufacturing industries and construction. Explored the casual link between CO2 emission and economic growth in 29 countries (14 developed and 15 developing). The study analyzed the connection between energy consumption of explanatory variables in developed and developing countries. Such EKC tested for historical perspective along with fuel prices and growth in Sweden in period of 1870-1997[2]. Explored the energy consumption and study of the electricity in Saudi Arabia with Time-Varying parameters vector autoregressive (TVP-VAR) in the period of 1970-2010. [3]. Study about the dynamic impact and economic output and Carbon emission from 1991-2012[4]. Tested the EKC hypothesis for solid waste generation with panel data from 1997-2010 in 32 European states [5]. Studied the technological progress and EKC, associated with economic growth and CO2 emission in panel data in 24 European nations from 1990-2013 [6]. Explored the transport energy by using EKC with hypothesis in EU-27 countries from 1995-2009 [7]. The main feature of this paper is to distinguish from other on the bases on research sample and some explanatory variable in developed and developing countries are creating effects on CO2 emission on other developed countries and how the manufacturing industries and military expenditure effects on the CO2 emission. The following section, logical structure and literature review highlighted the EKC hypothesis along with the relationship between CO2 emission and economic growth. In section 3, the data used for analysis along with econometric frameworks. Discussions and the empirical results are shown in section 4, while the final section of the paper concludes and provides implication policies with recommendations.

## 2. Literature review

A catholic part of specific literature explores the association between EKC and the national income of the countries, and greater environmental quality and their effects on developed and developing countries Table-1. According to Kuznets’ inverted U-hypothesis, initial stage as per capita national income of countries rise, inequality in income distribution rises after reaching the highest degree, where the country develops and its per capita income automatically rises in maximum level, and it falls as GDP per capita increases further [8]. Explored the study of 1955, and calculated the Kuznets’ ratio and found that, whereas developed countries tend to have lower degree of inequality, the developing countries tend to have a higher degree of inequality. That the evidence of inverted U-hypothesis, regarding the relationship between economic growth and inequality. It means that income inequalities where higher in developing countries compare to developed countries, but after that in particular stage, increase in economic growth will reduce the environmental pressure. In Table-3 summarized the turning points to identified the earlier studies.

**Table 1:**
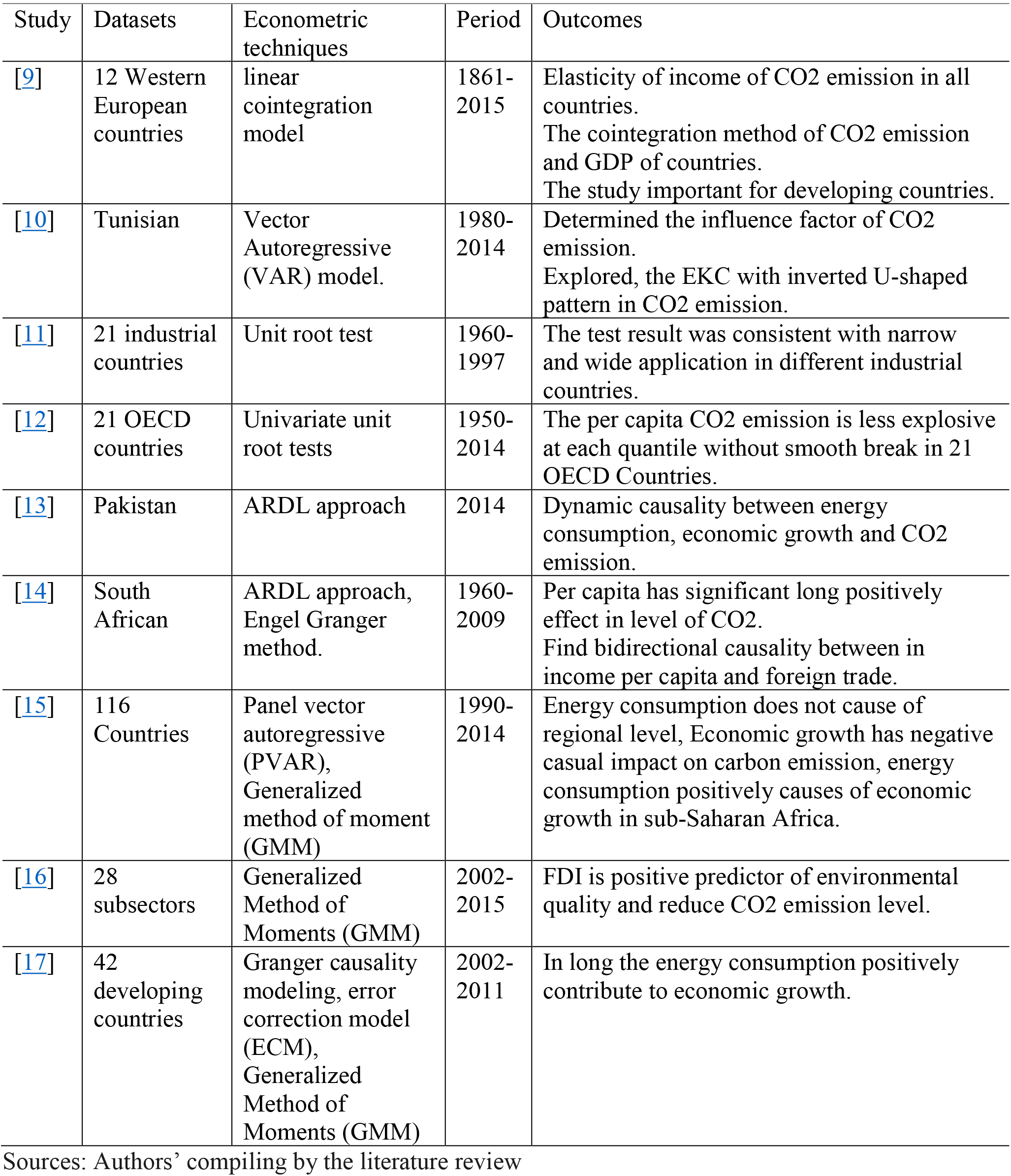
literature review of Economic growth and CO2 emission

| Study | Datasets | Econometric techniques | Period | Outcomes |
| --- | --- | --- | --- | --- |
| [9] | 12 Western European countries | linear cointegration model | 1861-2015 | Elasticity of income of CO2 emission in all countries.<br>The cointegration method of CO2 emission and GDP of countries.<br>The study important for developing countries. |
| [10] | Tunisian | Vector Autoregressive (VAR) model. | 1980-2014 | Determined the influence factor of CO2 emission.<br>Explored, the EKC with inverted U-shaped pattern in CO2 emission. |
| [11] | 21 industrial countries | Unit root test | 1960-1997 | The test result was consistent with narrow and wide application in different industrial countries. |
| [12] | 21 OECD countries | Univariate unit root tests | 1950-2014 | The per capita CO2 emission is less explosive at each quantile without smooth break in 21 OECD Countries. |
| [13] | Pakistan | ARDL approach | 2014 | Dynamic causality between energy consumption, economic growth and CO2 emission. |
| [14] | South African | ARDL approach, Engel Granger method. | 1960-2009 | Per capita has significant long positively effect in level of CO2.<br>Find bidirectional causality between in income per capita and foreign trade. |
| [15] | 116 Countries | Panel vector autoregressive (PVAR), Generalized method of moment (GMM) | 1990-2014 | Energy consumption does not cause of regional level, Economic growth has negative casual impact on carbon emission, energy consumption positively causes of economic growth in sub-Saharan Africa. |
| [16] | 28 subsectors | Generalized Method of Moments (GMM) | 2002-2015 | FDI is positive predictor of environmental quality and reduce CO2 emission level. |
| [17] | 42 developing countries | Granger causality modeling, error correction model (ECM), Generalized Method of Moments (GMM) | 2002-2011 | In long the energy consumption positively contribute to economic growth. |

Sources: Authors' compiling by the literature review

**Table 3:** Turning points reached earlier studies by pollutant type.

| Pollutant types | Study | Datasets | Period | Econometric techniques | Turning points |
| --- | --- | --- | --- | --- | --- |
| CO2 emission | [18] | 173 countries | 1990-2014 | Error correction model | (402,125.361 US\$) |
| CO2 emission | [19] | 20 countries | 1870-2014 | Bivariate model | \$18,955 and \$89,540 (in 1990 US\$) |
| CO2 emission | [20] | 128 countries | 1990-2014 | cross-sectional dependence and slope homogeneity tests | Significant |
| CO2 emission | [23] | 141 countries | 1970-2014 | Spatial Green Solow model | Statistically significant |
| CO2 emission | [24] | India | 1970-2015 | autoregressive distributed lag (ARDL) | USD 2937.77 |
| Renewable energy | [25] | Pakistan | 1970-2014 | autoregressive distributed lag (ARDL) | Significant |
| CO2 emission | [26] | 27 Chinese cities | 2001-2005 | Panel data parameter estimation | 34,328 CNY and 47,669 CNY |
| Industrial CO2 emission | [21] | USA | 1973-2015 | multilevel mixed-effect | Significant |
| CO2 emission | [27] | China | 1995-2011 | Input-output analysis | Significant |
| Fuel energy consumption | [28] | East Asian and Pacific countries | 1990-2014 | Generalized Method of Moment (GMM) | \$5112.65 |

Sources: Authors' compiling by the literature review

The EKC point starting from [18] showed that there is an inverted U-Shaped and relationship between per capita income and energy intensity in 173 countries and found CO2 emission by error correction model [19]. Explored the EKC hypothesis for panel of 20 countries with traditional inverted U-shaped relationship. [20] That study empirically related with economic and population growth and CO2 emission from 1990 to 2014. The cross-sectional study result dependent on slop homogeneity and heterogeneity. The common correlated effect means group (CCEMG), indicated the population size, economic growth and their significantly influence on the level of CO2 emission.

## 3. Data and Methodology

### Sample and variables

The data sample covers the period of 1977-2014 for panel consisting of the 29 (14 developed and 15 developing) countries. Table-2 indicate the variables, used for analysis, as well as their definitions and the sources of data are presented with different abbreviations. A part of preceding studies the EKC have already treated with different variables, like consumption of energy and economic growth. [9, 13, 17, 19], while the other new variables such as corruption, electricity consumption, population urbanization, industrial revolution provides more consideration [11, 21]. In CESFC, CEGFC, CELFC, CE, CEMIC control the trend of explanatory variables of AIT, CSE, IGD, CR, IF, ME and AL as well, high technology manufacturing sector includes high skill labor contribution in development and creating the significant effects on economy.

**Table 2:** Variables description for the analysis

| Variables | Definition | Unit measurement | Time frame availability | Data sources |
| --- | --- | --- | --- | --- |
| GDP | GDP per capita | Constant 2010 US dollars | 1977-2017 | World Bank (NY.GDP.MKTP. KD) |
| GDPC | GDP Per capita growth | Annual % | 1977-2017 | World Bank (NY.GDP.PCAP.KD. ZG) |
| CESFC | Co emissions from solid fuel consumption | kt | 1977-2014 | World Bank (EN.ATM.CO2E.SF.K T) |
| CEGFC | Co emissions from gaseous fuel consumption | kt | 1977-2014 | World Bank (EN.ATM.CO2E.GF.K T) |
| CELFC | Co emissions from liquid fuel consumption | kt | 1977-2014 | World Bank (EN.ATM.CO2E.LF.K T) |
| CE | Co emissions | kt | 1977-2014 | World Bank (EN.ATM.CO2E. KT) |
| CEMIC | Co emissions from manufacturing industries and construction | % of total fuel combustion | 1977-2014 | World Bank (EN.CO2.MANF. ZS) |
| ME | Merchandise Export | % of total merchandise exports | 1977-2016 | World Bank (TX.VAL.MRCH. R2. ZS) |
| AET | Arms export trend indicator | Value | 1977-2017 | World Bank (MS.MIL.XPRT. KD) |
| MI | Merchandise Import | % of total merchandise imports | 1977-2016 | World Bank (TM.VAL.MRCH. R2. ZS) |
| AIT | Arms import trend indicator | Value | 1977-2017 | World Bank (MS.MIL.MPRT. KD) |
| CSE | Commercial service export | Current US dollar | 1977-2017 | World Bank (TX.VAL.SERV.CD. WT) |
| IGD | inflation GDP deflator | Annual % | 1977-2017 | World Bank (NY.GDP.DEFL.KD. ZG) |
| CR | Coal rents (GDP) | % of GDP | 1977-2016 | World Bank (NY.GDP.COAL. RT. ZS) |
| IF | Insurance and financial service | % of commercial service exports | 1977-2017 | World Bank (TX.VAL.INSF.ZS. WT) |
| MIE | Military expenditure | % of GDP | 1977-2017 | World Bank (MS.MIL.XPND.GD.Z S) |
| AL | Agriculture land | % of land area | 1977-2015 | World Bank (AG.LND. AGRI. ZS) |

Sources: Selection based on databases' availability

While MI and AET control the GDP, high manufacturing and export development creating negative aspects. Initially per capita increase the wealth also increases the CO2 emission. However, arms import has created also significant effects on CESFC, CEGFC and CEMIC but not creating effects on CE Table-4. In empirical methodology, in what we follow, we start by testing unit roots all explanatory variables individually in panel data. If the variables have found non-stationary, we investigate the prevailing long run cointegration relationship and investigate their magnitude for long run stationary. We employ a class of panel unit root test and panel cointegration test individually on all explanatory variables, which allow the serial correlation among the cross section, i.e. the so-called second-generation test. Augmented IPS used by cross sectional [22] panel unit root test by Pesaran (2007) and as for panel cointegration used error-correction by Westerlund (2007), which both account for possible cross sectional dependencies for individual explanatory variables. Table-7 shows unit root test on level and Table-8 with first deference. The key variables-CO2 emission of GDP (Constant 2010 US dollars) and per capita GDP (Annual %) growth along with other explanatory Table-2, variables - in for both level and first difference. In the level case, we are unable to reject the null hypothesis, except for the GDP per capita growth, CO2 emission, arm import trend, commercial service export and inflation GDP deflator.

**Table 4:**
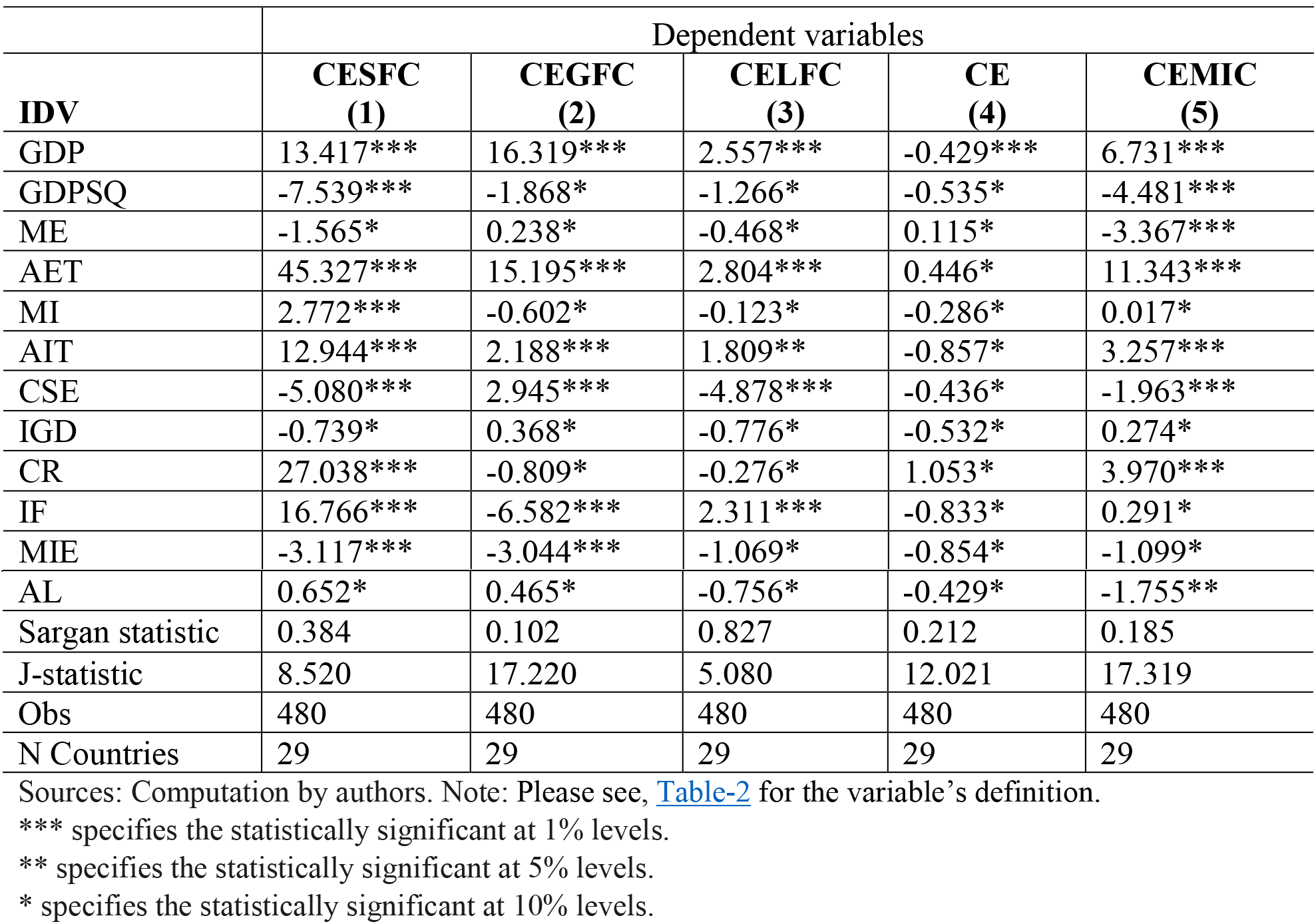
GMM regression with AB in n-Step

### 3.1 Econometric methods

EKC hypothesis, we followed the approach of [15, 17–19, 24–26, 29]. The long run relationship between polluted emission, GDP per capita, merchandise export, arms export, merchandise import, commercial service export, inflation GDP, coal rent, insurance and financial service, military expenditure and agriculture land, is given as follows:

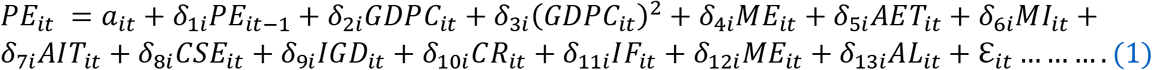

Where PE shows the polluted emission and i=1,….,29 and t=1977,….,2014 reveal the country and time, respectively whereas emission, which we take from solid, gases, and liquid fuel, CO2 emission and CO2 emission from manufacturing industries and construction. *a_it_* indicates the country fixed effect. The *δ*_1*i*_–*δ*_13*i*_ are parameters of long-run elasticities, which are related to each explanatory variable of the panel ε*_it_*, indicate estimated residuals, characterized for long-run equilibrium. Since the inverted U-Shaped EKC hypothesis ε_2*t*_, is expected to be positive and ε_3*t*_ is expected to be negative, also the monitoring value representing the turning points which is computed by *τ* = exp[−*β*_1_]/(2*β*_2_)] [19, 24, 29]. Additionally, the research aims to establish the casual link between manufacturing industries and construction, economic growth, arms export, commercial service export and coal rent (GDP). Additionally, the Generalized Method of Moments (GMM) yields steady and efficient parameter estimate in a regression, the explanatory variables are not strictly exogenous, heteroscedasticity and autocorrelation within exist [30]. The GMM is more efficient and effectual with an additional assumption that is the first difference of explanatory variables, which is turn allows the inclusion of more instruments. The GMM applied on 29 countries over 1977-2014 in order to analyze the impact of different explanatory variables on CO2 emissions. [31]. Thus, according to [32–34] first generation test such as common root-Levin, Lin (LLC), Chu and Breitung, individual Im, Pesaran, shin (IPS), Augmented Dickey-Fuller (ADF), and individual root-Fisher-PP, and Hadri have been computed individually from all explanatory variables. Afterward, we computed the cointegration test by Pedroni, [35] Kao from Engle-Granger based and Fisher [36] (combined Johansen).

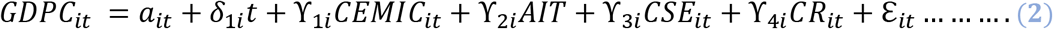

Where i=1,….,29 and t=1977,….,2014 for each country in panel data. Besides, the parameters *a_i_* and *δ_i_* indicate the fixed effect and deterministic trend. It is computing by Engle-Granger, long term model, specified in Eq (2) is estimated in which one period lagged and residual as error correction term.

The dynamic error correction model is represented below:

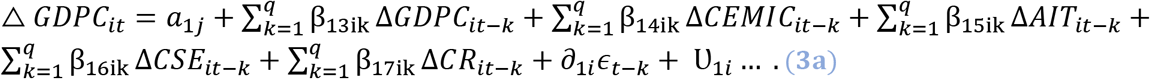

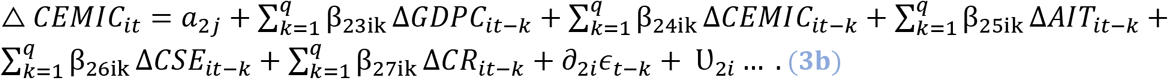

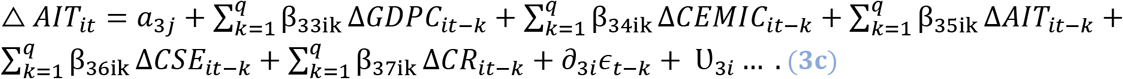

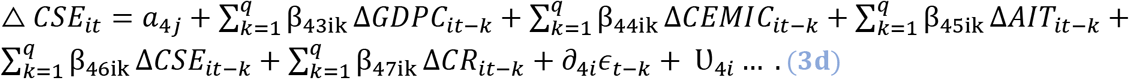

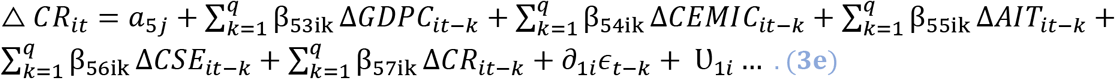

Where the first-difference operator indicates by Δ, the lag of length specified by *q* at one according to likehood ratio test, and 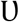 specify serial uncorrelated error term.

## 4. Results

### 4.1 Descriptive statistics, correlation and unit root examination

Table-5 shows the descriptive statistics of the particular variables of high mean value over the period of 1977-2014, countries by the type of pollutant emissions, China (CESFC, CE), USA (CEGFC, CELFC, AET, CSE) and India (AIT) show the highest mean value Fig-3 Although, Morocco (CR), Mexico (MI, ME), Philippine and Canada (ME), Mexico and Panama (MIE), Costa Rica and Argentina (IF) register the lowest mean value. Table-6 indicate the term of matrix correlation, relationships between energy consumption and selected instrumental variables, emission such as CESFC, CEGFC, CELFC, CE and CEMIC were noticed. Fig-4 explored the value of mean, the manufacturing industries and construction increase continuously comparatively solid, liquid and gaseous fuel consumption. The result computed by GMM method and in order to remove inconvenience, consider stationary test according to cross section independence in first generation unit root test in common root and individual intercept in level and 1^st^ generation Table-7. [37–39] As we notice the variables are non-stationary in their level, and become stationary after 1^st^ difference Table-8.

**Fig-3:**
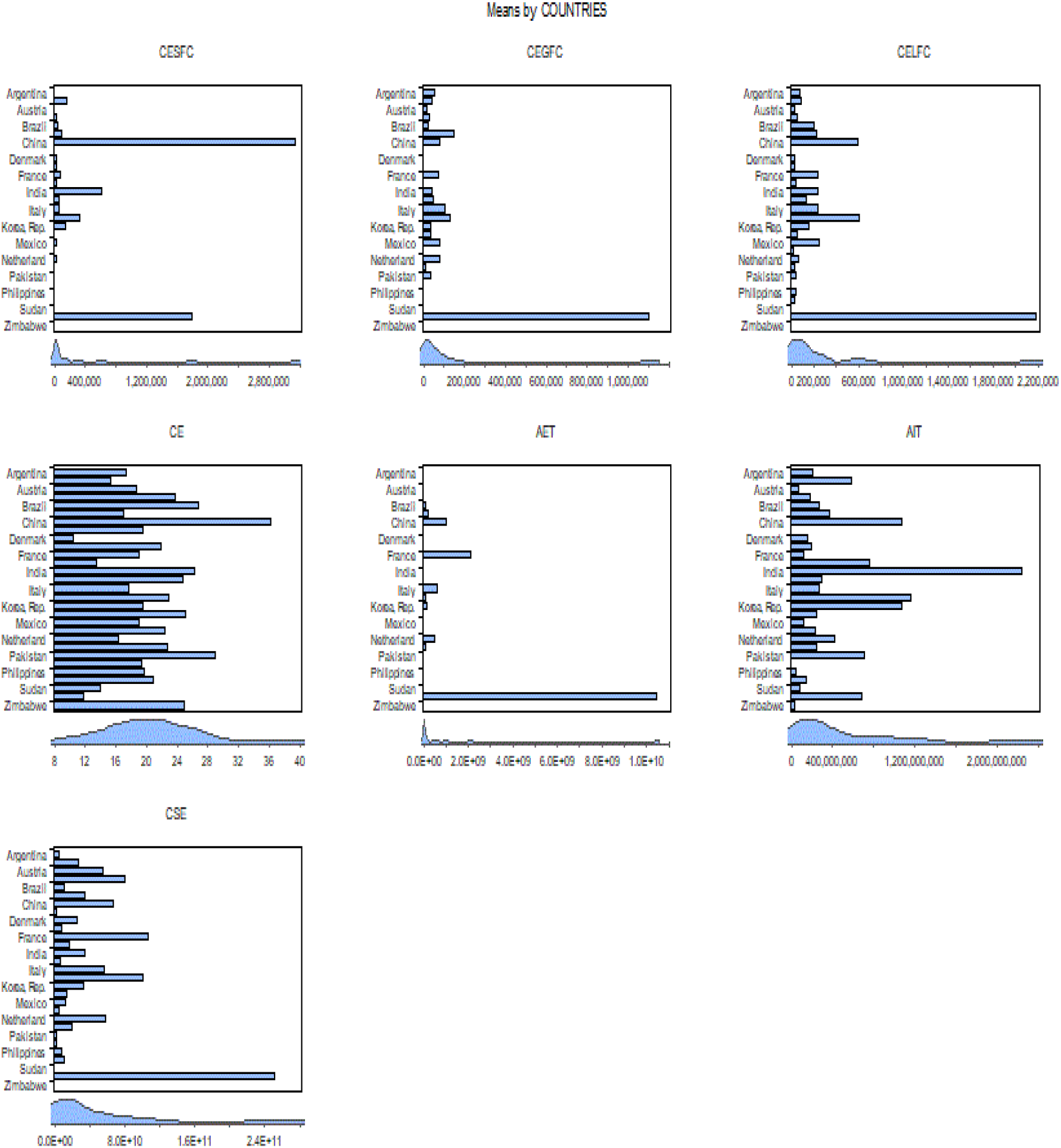
Highest mean valuation of pollutant emission by 29 countries

**Fig-4:**
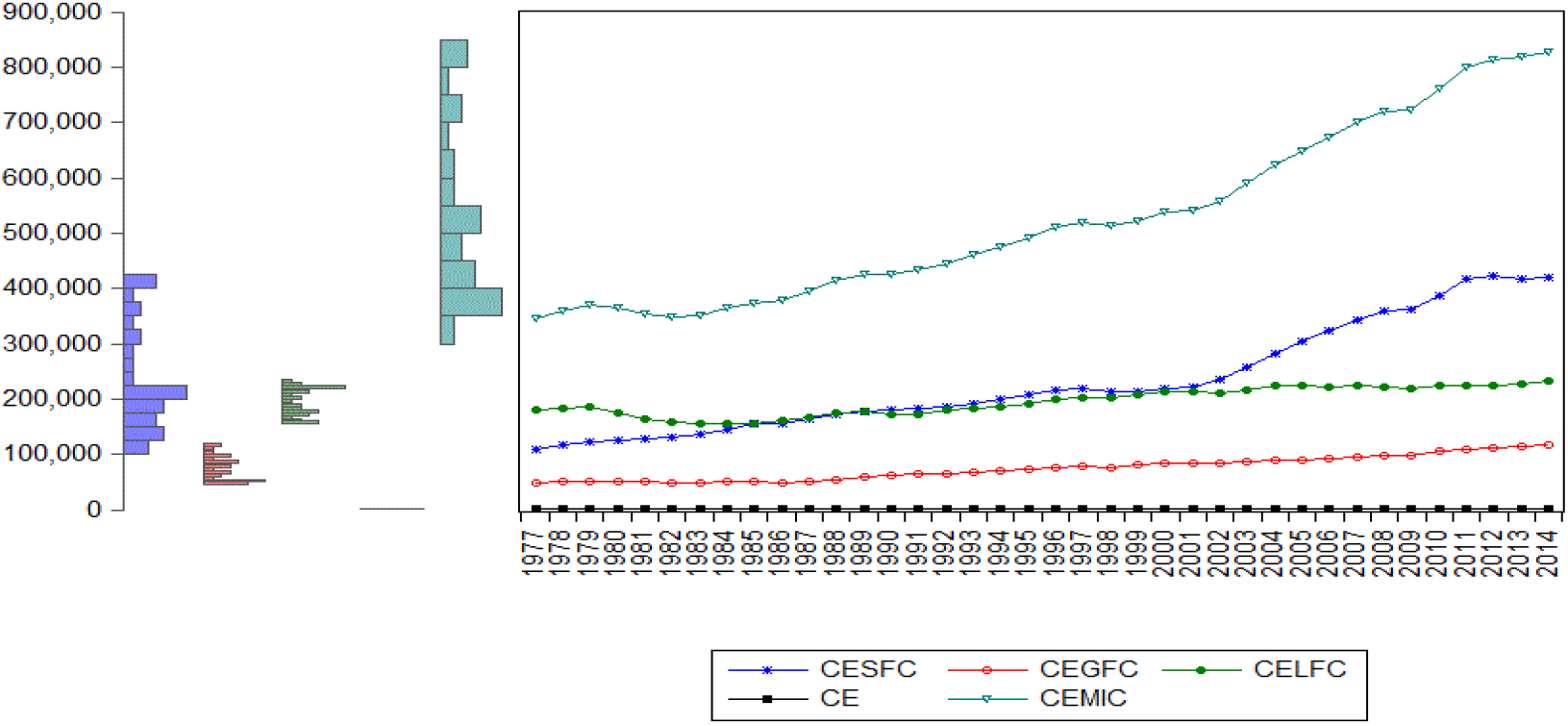
Mean value of pollutant emissions by years

**Table 5:** Descriptive statistics (Raw data)

| Variables | Mean | Median | Max | Min | Sta.Dev. | Skewness | Kurtosis | Jarque-Bera | Prob | Obs |
| --- | --- | --- | --- | --- | --- | --- | --- | --- | --- | --- |
| GDP | 1,080,000 m | 309,000, m | 16,200,000, m | 6,750, m | 2,250,000m | 4.11 | 22.03 | 19,728.63 | 0.00 | 1,102 |
| GDPC | 2.209713 | 2.26 | 13.64 | -15.32 | 3.73 | -0.73 | 6.46 | 646.47 | 0.00 | 1,102 |
| CESFC | 231,943.80 | 21,536.29 | 7,499,587.00 | -113.68 | 752,749.80 | 5.98 | 47.32 | 96,758.81 | 0.00 | 1,102 |
| CEGFC | 74,688.62 | 17,552.10 | 1,432,767.00 | 0.00 | 202,645.60 | 4.84 | 26.34 | 29,322.15 | 0.00 | 1,102 |
| CELFC | 195,177.90 | 56,612.98 | 2,494,601.00 | 1,452.13 | 411,692.50 | 4.04 | 19.65 | 15,727.17 | 0.00 | 1,102 |
| CE | 525,318.10 | 114,734.90 | 10,291,927.00 | 2,002.18 | 1,276,136.00 | 4.23 | 22.95 | 21,557.75 | 0.00 | 1,102 |
| CEMIC | 20.54 | 19.53 | 49.15 | 0.00 | 7.29 | 0.69 | 3.77 | 115.22 | 0.00 | 1,102 |
| ME | 2.15 | 1.08 | 28.83 | 0.00 | 3.55 | 4.25 | 24.91 | 24,815.98 | 0.00 | 1,079 |
| AET | 943 m | 76 m | 15,700 m | 0.00 | 2,610 m | 3.79 | 17.03 | 6,592.15 | 0.00 | 622 |
| MI | 2.13 | 0.96 | 27.10 | 0.00 | 3.23 | 3.31 | 17.06 | 10,832.16 | 0.00 | 1,077 |
| AIT | 444 m | 200 m | 5,320, m | 0.00 | 638 m | 2.88 | 13.85 | 6,421.31 | 0.00 | 1,022 |
| CSE | 34,300 m | 10,100 m | 721,000 m | 13.5 m | 70,000 m | 5.13 | 38.34 | 55,969.20 | 0.00 | 992 |
| IGD | 25.76 | 4.61 | 3,057.63 | -27.05 | 176.35 | 13.00 | 186.77 | 1,581,752.00 | 0.00 | 1,102 |
| CR | 0.22 | 0.00 | 8.71 | 0.00 | 0.66 | 6.18 | 58.31 | 147,507.50 | 0.00 | 1,102 |
| IF | 3.56 | 2.30 | 22.08 | -2.28 | 3.66 | 1.36 | 4.88 | 439.85 | 0.00 | 964 |
| MIE | 2.39 | 2.12 | 10.67 | 0.00 | 1.47 | 1.04 | 4.67 | 316.74 | 0.00 | 1,071 |
| AL | 40.62 | 44.82 | 71.54 | 2.46 | 18.69 | -0.43 | 2.17 | 64.52 | 0.00 | 1,079 |

Note: m indicates million. Sources: Definition of variable available in Table-2

**Table 6:**
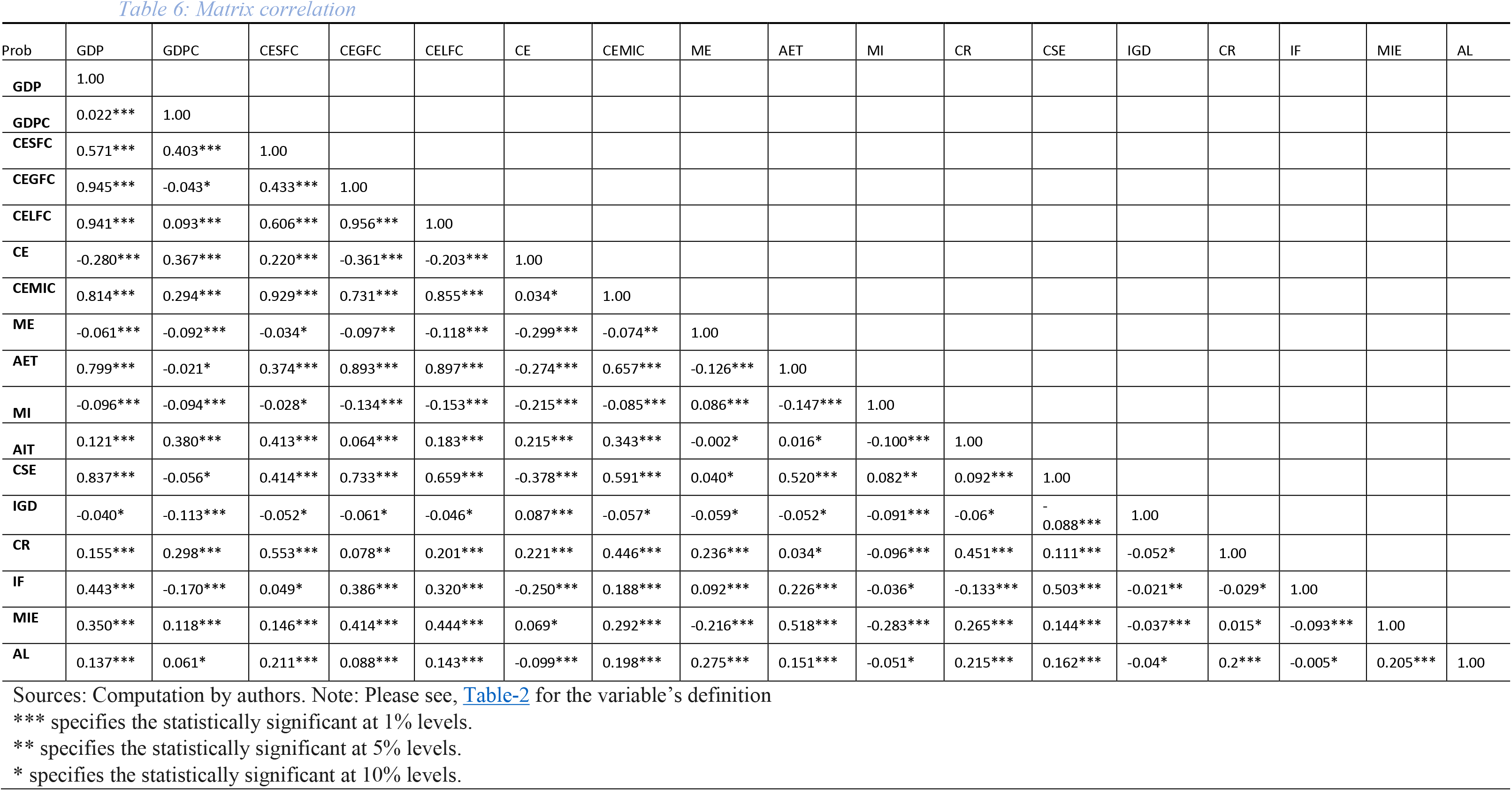
Matrix correlation

**Table 7:** Unit root of individual variables (Level)

| Level |  |  |  |  |  |  |  |  |  |  |  |
| --- | --- | --- | --- | --- | --- | --- | --- | --- | --- | --- | --- |
| Variables | Individual intercept |  |  |  |  | Individual intercept and trend |  |  |  |  |  |
|  | CR | Individual root |  |  | Hadri | CR |  | Individual root |  |  | Hadri |
|  | LLC | IPS | ADF | PP |  | LLC | Breitung | IPS | ADF | PP |  |
| GDP | 8.739 | 13.65 | 16.039 | 17.32 | 19.797*** | 3.537** | 6.503 | 5.835 | 31.959** | 40.602** | 15.174*** |
| GDPC | -13.66*** | -12.975*** | 280.248*** | 380.343*** | 4.285*** | -13.69*** | -14.14*** | -12.784*** | 263.21*** | 425.104*** | 2.308*** |
| CESFC | 3.314** | 3.172** | 49.643** | 51.126** | 17.197*** | 1.202** | 6.941 | 2.330** | 60.639** | 64.380** | 13.552*** |
| CEGFC | 3.025** | 6.834 | 18.342 | 25.46** | 16.917*** | 4.958 | 7.018 | 5.049 | 41.332** | 44.202** | 7.476*** |
| CELFC | 2.695** | 3.300** | 59.771** | 55.412** | 16.591*** | -1.041** | 2.297** | 1.086** | 59.143** | 36.310** | 9.302*** |
| CE | -4.601*** | -1.436** | 74.517** | 89.044*** | 16.995*** | -0.633** | -0.7237*** | -0.783** | 67.931** | 83.414*** | 11.111*** |
| CEMIC | 3.992*** | 6.072 | 26.559** | 28.959** | 17.839*** | 2.607*** | 6.2 | 3.839** | 36.388** | 39.174** | 14.375*** |
| ME | 1.475** | 2.156** | 42.353** | 68.413** | 18.215*** | -1.518** | -1.482** | -2.352** | 81.068** | 102.354*** | 7.125*** |
| AET | -2.149*** | -3.04*** | 79.523 | 110.819 | 11.871*** | .717** | -5.135*** | -1.167*** | 61.545 | 87.606 | 3.010*** |
| MI | 3.707** | 5.879 | 37.712** | 52.798** | 18.826*** | -2.416** | 2.854** | -2.118** | 88.14** | 113.464*** | 13.400*** |
| AIT | -6.969*** | -8.674*** | 185.301*** | 248.052*** | 6.569*** | -6.696*** | -5.215*** | -6.628*** | 139.403*** | 203.930*** | 7.207*** |
| CSE | 11.033 | 14.16 | 3.186 | 1.533 | 19.175 | 2.628** | 6.902 | 5.625 | 19.432 | 15.76 | 14.747*** |
| IGD | -5.321*** | -6.227*** | 147.989*** | 202.607*** | 2.050** | -6.39*** | -5.44*** | -6.066*** | 144.463*** | 204.946*** | 7.540*** |
| CR | -3.471*** | -3.471*** | 72.654*** | 117.598*** | 4.397*** | -3.677*** | -3.956*** | -2.407 | 61.733** | 81.987*** | 9.117*** |
| IF | -2.095*** | -2.917*** | 96.901*** | 113.746*** | 11.374*** | -6.89*** | -3.299*** | -4.257*** | 107.359*** | 127.867*** | 127.867*** |
| MIE | -3.048** | -0.505** | 62.802** | 60.849** | 15.849*** | -1.574*** | -1.968*** | -0.695 | 60.823 | 68.363 | 8.344*** |
| AL | -1.654** | 3.599** | 39.238** | 53.892** | 14.337*** | -0.876** | 1.459** | 1.960** | 38.680** | 45.758** | 12.985** |

Source: Computation by authors. Note: Please see, Table-2 for the variable's definition

**Table 8:** Unit root of individual variables (First difference)

| First difference |  |  |  |  |  |  |  |  |  |  |  |
| --- | --- | --- | --- | --- | --- | --- | --- | --- | --- | --- | --- |
| Variables | Individual intercept |  |  |  |  | Individual intercept and trend |  |  |  |  |  |
|  | CR | Individual root |  |  | Hadri | CR |  | Individual root |  |  | Hadri |
|  | LLC | IPS | ADF | PP |  | LLC | Breitung | IPS | ADF | PP |  |
| GDP | -6.892*** | -9.509*** | 217.311*** | 321.655*** | 14.3024*** | -11.7028*** | -7.733*** | -11.631*** | 229.986*** | 379.76*** | 9.978*** |
| GDPC | -26.19*** | -29.252*** | 692.647*** | 771.738*** | -5.267** | -23.208*** | -15.864*** | -26.604*** | 686.887*** | 5352.97*** | 9.362*** |
| CESFC | -12.81*** | -18.857*** | 430.863*** | 650.227*** | 7.574*** | -11.713*** | -6.668*** | -18.948*** | 431.265*** | 970.338*** | 2.286** |
| CEGFC | -8.981*** | -11.885*** | 293.098*** | 593.465*** | 4.065*** | -10.029*** | -2.635*** | -11.885*** | 243.819*** | 1059.95*** | 3.771*** |
| CELFC | -11.43*** | -14.09*** | 311.425*** | 584.042*** | 2.304*** | -10.102** | -7.062*** | -12.884*** | 269.772*** | 1016.83*** | 7.214*** |
| CE | -15.56*** | -20.566*** | 474.701*** | 779.588*** | .8016** | -13.235** | -13.79*** | -19.213*** | 429.375*** | 1259.51*** | 4.450*** |
| CEMIC | -9.513*** | -14.738*** | 334.107*** | 65.894*** | 10.289*** | -8.196*** | -5.221*** | -13.383*** | 281.423*** | 688.814*** | 3.978*** |
| ME | -14.77** | -20.001** | 462.271*** | 793.076*** | -0.17** | -12.353*** | -10.094** | -18.622*** | 389.420*** | 1483.56*** | 4.355*** |
| AET | -6.224*** | -11.817*** | 226.406*** | 488.133*** | 5.059*** | -1.429** | -4.816*** | -7.762*** | 175.094*** | 906.84*** | 25.403*** |
| MI | -11.5*** | -20.324*** | 468.653*** | 745.406*** | 2.193** | -7.822*** | -4.919*** | -18.521*** | 410.287*** | 2596.82*** | 5.858*** |
| AIT | -19.22*** | -22.654*** | 520.978*** | 782.556*** | -1.269*** | -15.856*** | -8.849*** | -17.968*** | 426.592*** | 3790.04*** | 4.138*** |
| CSE | -10.58*** | -14.029*** | 318.835*** | 551.193*** | 12.867*** | -11.013*** | -8.543*** | -13.078*** | 330.713*** | 863.553*** | 5.204*** |
| IGD | -22*** | -24.622*** | 589.251*** | 790.052*** | 3.526*** | -19.072*** | -14.156*** | -22.111*** | 486.624*** | 3147.26*** | 23.308*** |
| CR | -21.31*** | -25.57*** | 552.546*** | 653.620*** | -2.768*** | -18.106*** | -15.786*** | -23.328*** | 468.263*** | 656.767*** | 1.455** |
| IF | -22.4*** | -19.626*** | 442.645*** | 740.570*** | .0.187** | -29.65*** | -11.275*** | -16.375*** | 361.590*** | 1509.40*** | 6.791*** |
| MIE | -12.71*** | -14.969*** | 326.141*** | 635.900*** | 3.653*** | -10.897*** | -11.559*** | -12.816*** | 262.854*** | 1462.23*** | 16.187*** |
| AL | -8.498*** | -12.687*** | 291.138*** | 559.093*** | 5.3833*** | -7.563*** | -5.376*** | -10.829*** | 238.115*** | 796.856*** | 7.260*** |

Source: Computation by authors. Note: Please see, Table-2 for the variable's definition

### 4.2 Panel regression analysis

Table-4 indicate the GMM regression method with AB in n-step. In the GMM estimation, the explanatory variable individually estimated regression with dependent variables. The panel data study by providing the solution of common problems in different developed and developing countries; the heterogeneity of behavior of individual explanatory variable, the endogenous and simultaneity by bidirectional causality problem. This research paper will estimate a dynamic model (where the endogenous variables are included as explanatory variables along with more than one lag). The white period method applies for coefficient covariance method individually for computation of CESFC, CEGFC, CELFC, CE and CEMIC with other explanatory variables. The difference cross sectional period was used for cross section in none period, the GMM iterations was computing in 2-step, that varies by cross-section in white period.

According to Sargan statistic, all estimated models are statistically highly significant, and the value of J-Statistic, that could be explained between 5.08 and 17.31 of the variability in pollutant emission. Hence in the model, where the same number in instrument as a parameter, the optimized value of the objective function is zero. If the number of instruments increased than parameters, the optimized value will be greater than zero, and the J-statistic used as test of over-identifying moment condition. The J-statistics and instrumental rank, reported by Sargan statistics, where the instrumental rank greater in individual model, than number of estimated coefficients, we may use to construct Sargan test over the identifying restrictions. While in the null hypothesis overidentifying restriction are valid, the J-statistic in panel equation is different from the ordinary equation, where the Sargan statistics is distributed as a *χ*(*ρ – k*). Where the estimated coefficient is k and instrumental rank is *ρ* individual in each model. The Sargan test was computed in CESFC by scalar pval = @chisq (8.50,9.0) individually. The related coefficient of GDP per capita and squared GDP per capita are statistically significant in all estimated model, except model 4, the EKC hypothesis is confirmed in case of CE negatively impact. Furthermore, estimated regression appears to fit the data by the value of Sargan test, they can explain all most 10% to 82% of the pollutant emission. The inverted U-Shaped curve emerges in all cases of harming secretions, except CE, with regard of GDPSQ, MI, AIT, CSE, IGD, IF, ME and AL; knowledge that expectation ecological damage reduction is not support positively in estimated models, show a negative influence on pollutant emission. Also, we notice with some exceptional the renewable energies consumption reduces the pollution emission, like the higher GDP implies higher production and more insurance and financial services acquired [40]. In the term of merchandise export (ME) like [41]. The results of the variables employed to control for the scale effect and pollution conditions.^13^

Fig-5 reveals the plotted graphs between GDP and pollutant emission. The EKC hypothesis appaired to be sustained since the inverted U-shaped curve tend to be fit properly in CESFC, and also indicated the sequence of U-shaped, in the term of CEGFC and CESFC, curve straightly going upward and we notice that the turning points are not in line. Hence in carbon emission the EKC curve coming down and notice that after high technology in industries and export reduce the level of EKC. In last CEMIC the intensity of emission continuously in developing countries. Furthermore, [42] specified a higher likelihood of identifying turning points in case of developed to developing countries.

**Fig-5:**
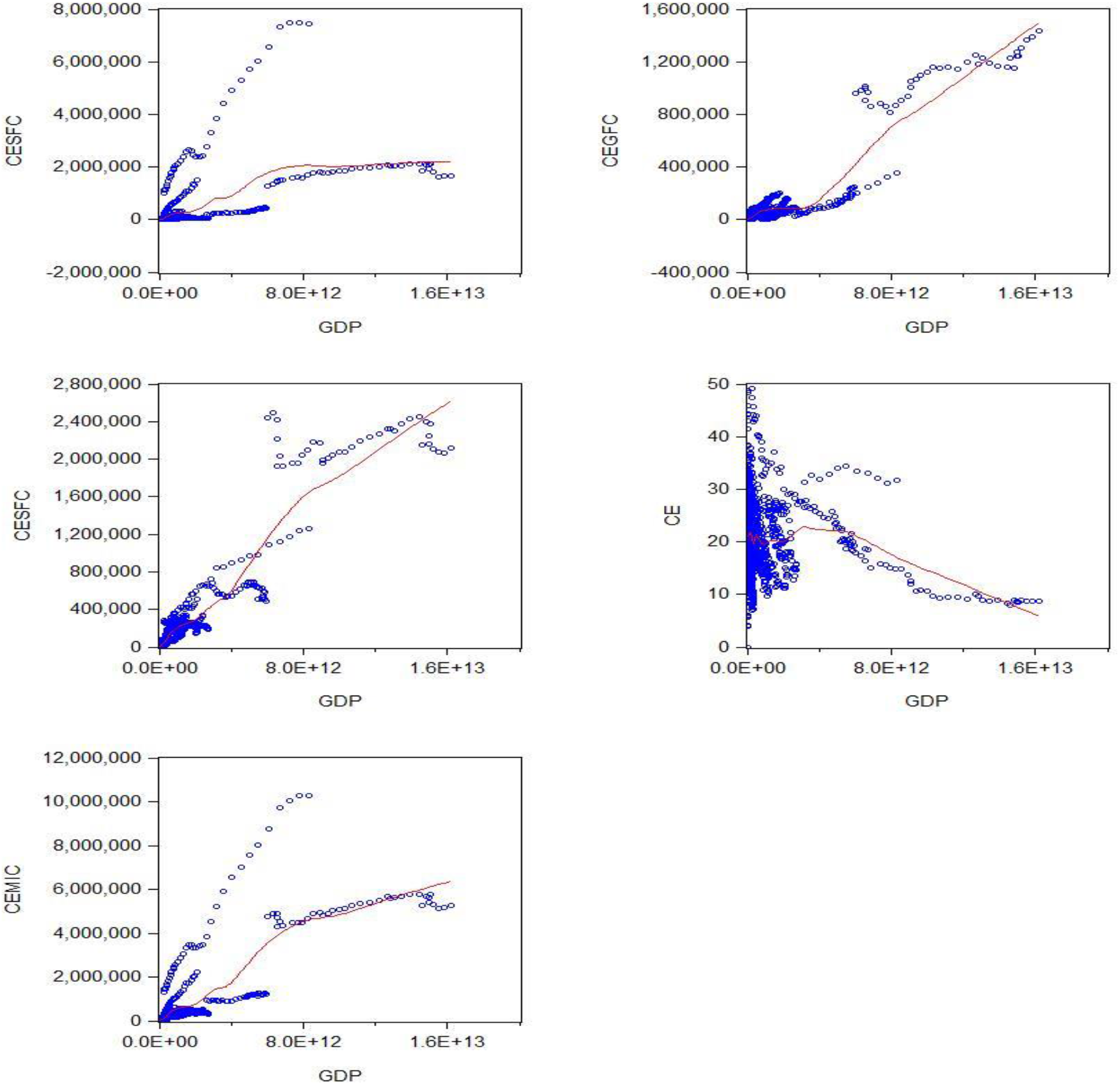
Plotted graph between GDP per capita and CESFC, CEGFC, CESFC, CE and CEMIC

### 4.3 Co-integration and causal investigation

In the co-integration, the Padroni panel test [35] is explored in Table-9. The dimensional approach of statistics, the autoregressive coefficient in the different developed and developing countries [32, 43] for the unit root test on the estimated residual consideration for heterogeneity across the country and time factor. And the analysis of long-run cointegration relationships has been taken from developed and developing countries in modern series analysis.

**Table 9:** Pedroni (Engle-Granger based) Test

| Panel A: Wintin-dimension |  |  |  |  |  |  |
| --- | --- | --- | --- | --- | --- | --- |
| Panel co-integration test | Individual intercept |  | Individual intercept and trend |  | No intercept or trend |  |
|  | Statistic | Weighted Statistic | Statistic | Weighted Statistic | Statistic | Weighted Statistic |
| Panel v-Statistic | 2.737*** | -3.115* | 22.167*** | -0.938* | -1.480* | -3.404* |
| Panel rho-Statistic | 0.658* | 1.524* | -1.226* | 2.162* | -0.248* | 1.393* |
| Panel PP-Statistic | -0.388* | 2.749* | -3.865*** | 0.195* | -0.144* | 1.822* |
| Panel ADF-Statistic | -0.242* | 2.993* | -3.652*** | 4.061 | -0.235* | 1.105* |
| Panel B: Between- dimension |  |  |  |  |  |  |
| Panel co-integration test | Individual intercept |  | Individual intercept and trend |  | No intercept or trend |  |
|  | Statistic |  | Statistic |  | Statistic |  |
| Group rho-Statistic | 3.275* |  | 4.086 |  | 2.660* |  |
| Group PP-Statistic | 1.776* |  | 0.617* |  | 1.635* |  |
| Group ADF-Statistic | 2.981* |  | 0.236* |  | 2.977* |  |

Source: Computation by authors. The lag length was selected by Schwarz Info criterion.
Note: Please see, Table-2 for the variable's definition
\*\*\* specifies the statistically significant at 1% levels.
\*\* specifies the statistically significant at 5% levels.
\* specifies the statistically significant at 10% levels.

Table-9 The Padroni panel test in panel A, ADF statistically reject the null hypothesis of no cointegration with individual intercept, trend and No intercept or trend. The statistically mean value of individual autoregressive coefficient related with unit root test of individual each developed and developing state. In the panel B, the co-integration employed with rho, PP and ADF statistics, and explored by the Kao Table-10 in Engle Granger based test, the ADF (t-statistics) is 2.490 (sig) with residual variance. Where the vector of co-integration is homogenous in different states. The result provides hypothesis of co-integration of developing and developed states variables.

**Table 10:** Kao (Engle Granger based) test

| ADF (t-Statistic) | Residual variance | HAC variance |
| --- | --- | --- |
| 2.490*** | 8.24E+21 | 2.64E+22 |

Source: Computation by authors. The lag length was selected by Schwarz Info criterion.
\*\*\* specifies the statistically significant at 1% levels.
\*\* specifies the statistically significant at 5% levels.
\* specifies the statistically significant at 10% levels.

The third test is a Fisher, that approach is used to underlying Johansen methodology by panel cointegration test [44], showed in Table-11. This panel co-integration test aggregates with p-value of individual Johansen trace statistics and eigen-value [45]; also reject the null hypothesis of no co-integration.

**Table 11:** Fisher (Combined Johansen) test

| Hypothesized No. of CE(s) | Fisher Stat.* (from trace test) | Fisher Stat.* (from max-eigen test) |
| --- | --- | --- |
| None | 135.8*** | 102.0*** |
| At most 1 | 64.86*** | 61.32*** |
| At most 2 | 32.51* | 32.51* |

Source: Computation by authors. The lag length was selected by Schwarz Info criterion and Probabilities are computed using asymptotic Chi-square distribution
\*\*\* specifies the statistically significant at 1% levels.
\*\* specifies the statistically significant at 5% levels.
\* specifies the statistically significant at 10% levels.

Onward, since the variable are co-integrated, a panel vector error correction model is estimated in order to perform Pairwise Granger Causality test Table-12, we reject the null that GDPC does not granger cause CEMIC, and also in the opposite direction.

**Table 12:** Pairwise Granger Causality Tests

| Null Hypothesis: | Obs | F-Statistic |
| --- | --- | --- |
| CEMIC does not Granger Cause GDPC | 1073 | 13.732*** |
| GDPC does not Granger Cause CEMIC |  | 47.520*** |
| AIT does not Granger Cause GDPC | 965 | 16.161*** |
| GDPC does not Granger Cause AIT |  | 4.293*** |
| CSE does not Granger Cause GDPC | 961 | 1.510 |
| GDPC does not Granger Cause CSE |  | 11.346*** |
| CR does not Granger Cause GDPC | 1073 | 21.069*** |
| GDPC does not Granger Cause CR |  | 5.530*** |
| AIT does not Granger Cause CEMIC | 965 | 56.007*** |
| CEMIC does not Granger Cause AIT |  | 6.348*** |
| CSE does not Granger Cause CEMIC | 961 | 133.750*** |
| CEMIC does not Granger Cause CSE |  | 22.872*** |
| CR does not Granger Cause CEMIC | 1073 | 51.272*** |
| CEMIC does not Granger Cause CR |  | 3.889*** |
| CSE does not Granger Cause AIT | 863 | 0.498 |
| AIT does not Granger Cause CSE |  | 3.675*** |
| CR does not Granger Cause AIT | 965 | 4.190*** |
| AIT does not Granger Cause CR |  | 3.319** |
| CR does not Granger Cause CSE | 961 | 0.009 |
| CSE does not Granger Cause CR |  | 0.929 |

Source: Computation by authors. The lag length was selected by Schwarz Info criterion.
Note: Please see, Table-2 for the variable's definition
\*\*\* specifies the statistically significant at 1% levels.
\*\* specifies the statistically significant at 5% levels.
\* specifies the statistically significant at 10% levels.

Table-13. indicate Vector Error Correction (VEC), with cointegration restrictions B (1,1) =1 and the convergence attained after 1 iteration with t-statistics and Standard error Fig-6 The specification of VEC has five (*k*=5) endogenous variables, GDPC, CEMIC, AIT, CSE and CR, the exogenous intercept C(d=1) and lags include 1 to 2 (*p*=1). Thus, there are (kp+d=6) regression of each of the three equation in the VEC individually.

**Table 13:** Vector Error Correction Model

| Error Correction: | Cointegration | Standard error | t-statistics | R-squared | F-statistic |
| --- | --- | --- | --- | --- | --- |
| D(GDPC) | -0.033 | -0.01641 | -2.04368 | 0.162128 | 26.05806 |
| D(CEMIC) | 2034.459 | -209.35 | 9.71799 | 0.690684 | 300.7023 |
| D(AIT) | 1093653 | -1525124 | 0.71709 | 0.049723 | 7.046426 |
| D(CSE) | 4.62E+08 | -3.20E+07 | 14.2740 | 0.291235 | 55.33518 |
| D(CR) | -0.002747 | -0.00132 | -2.08210 | 0.069086 | 9.994087 |

Source: Computation by authors. The lag length was selected by Schwarz Info criterion in cointegration restriction.
Note: Please see, Table-2 for the variable's definition
\*\*\* specifies the statistically significant at 1% levels.
\*\* specifies the statistically significant at 5% levels.
\* specifies the statistically significant at 10% levels.

**Fig-6:**
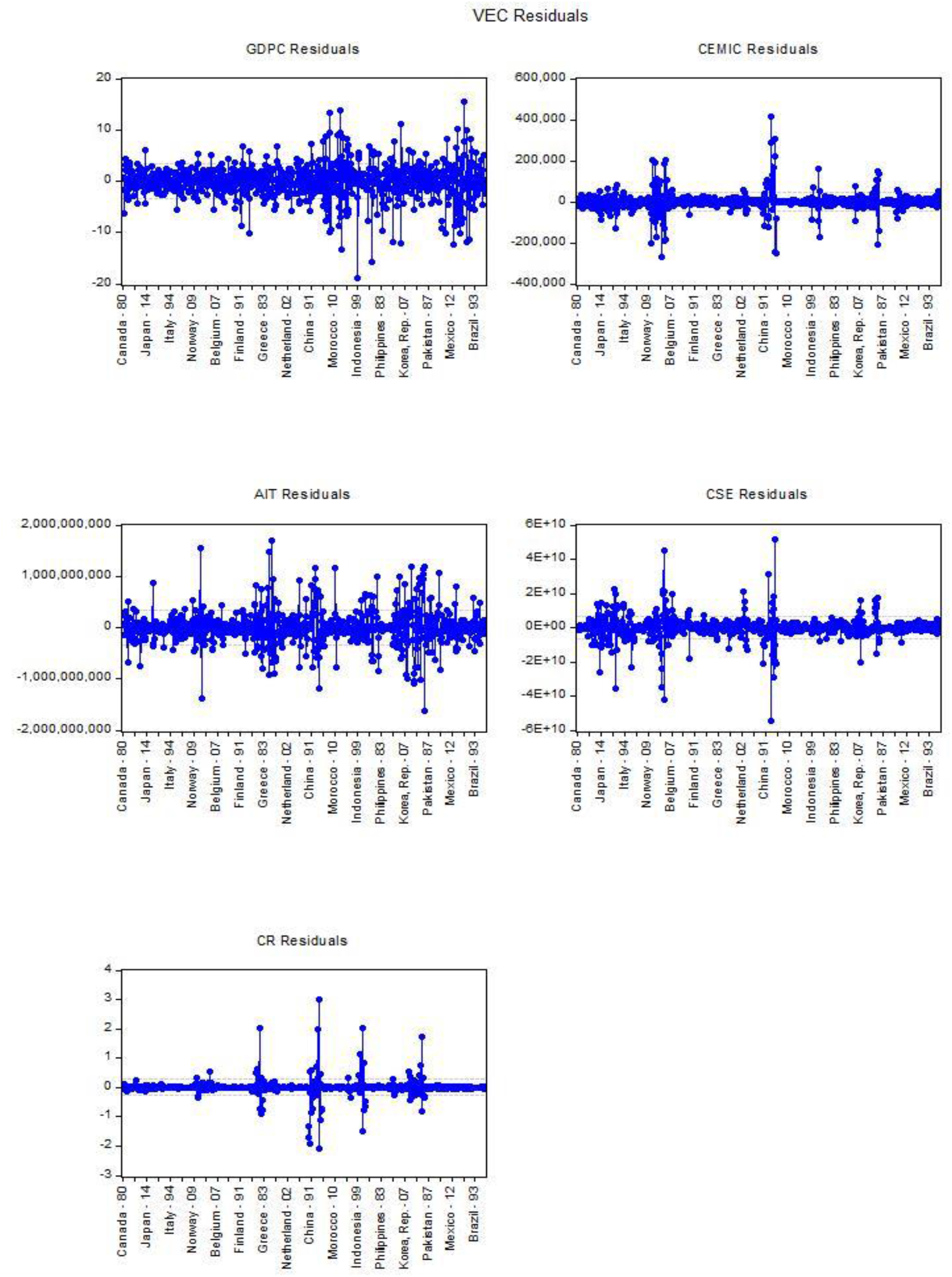
VEC Residuals by states

The effect of CEMIC has also been investigated by using impulse response by Cholesky one S. D (d.f. adjusted) innovation in decomposition method Fig-7, the impulse response of emission shock to Eq 3(a)–3(b) individually. The level of significant of impulse function has been investigated at 95%. The result from variance decomposition indicate the individual variables effects. In order to measure the deviation method, which impulse to GDPC are explained by CEMIC, AIT, CSE and CR. Eq (3a), according to VAR lag order selection criteria the endogenous variables indicated significant relationship in lag-2 at Schwarz information criteria (SC) and lag-17 at Hannan-Quinn information criteria, the CO2 emission is not too much efficient in lag-17, therefore the Johansen Fisher Panel Cointegration Test is applied in lag (2-1=1), it indicate the significant p-value (0.000) in model Table 13 are cointegrated in that case we use Vector Error Correction Estimates (VECM) in lag-1 with cointegration restrictions. The t-test in error correction model indicate significant relationship among GDP per capita and manufacturing industries and construction (CEMIC) with 9.718 which is more than 1.96, concerning Eq 3(b) identify that 69.0% manufacturing industries and construction has influence on the level of GDP per capita with F-statistics (300.702) comparatively others. Hence, the commercial service export (CSE) also indicate the significant relationship with GDP per capita in Eq 3(d), 29.123% has influence on the level of GDP per capita with F-statistics (55.335). In Eq 3(c) noticed the statistically insignificant influence on arms import (AIT) with 4.9% by GDP per capita. Eq 3(e) indicate the coal rent (CR) has not influence on GDP per capita with 6.90%. Moreover, the vector error correction term statistically significant in two endogenous variables, the analysis suggests that the above explanatory variables Table 13. are the main sources of volatility in different states by GDP per capita.

**Fig-7:**
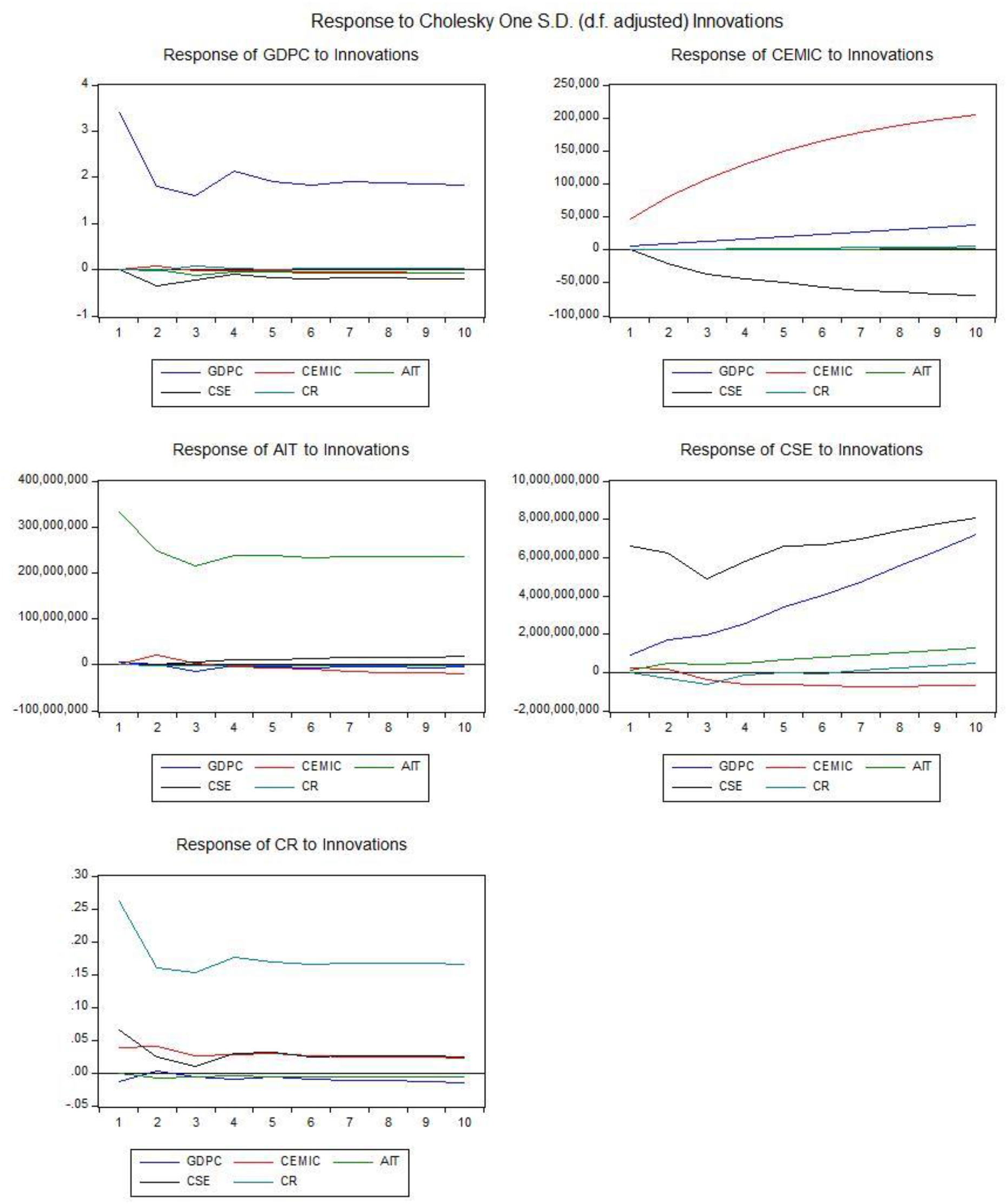
Impulse response

## 5. Conclusion

The objective of this research study was to determine the EKC hypothesis and afterward the causal relationships between carbon emission solid, liquid and gases fuel, merchandise export, economic growth, arms export trend, coal rents and military expenditure, for a panel consisting of 29 countries the period 1977-2014.

In the panel data, we noticed cross-sectional dependence in each of the variables, we employed Generalized Method of Movement/Dynamic Panel data, transformation of first deference with white period instrumental weighted mix. The results of GMM regression confirmed the acquired hypothesis for emission of CO2 emission from liquid fuel consumption, CO2 emission from manufacturing industries, where the outcome of GMM estimation corroborated, furthermore the EKC approach for solid, liquid and fuel consumption emission and CO2 emission.

Moreover, the estimation of GDP per capita with a panel vector error correction model in order to performed Pairwise Granger Causality test. The model shows a short run unidirectional causality from GDP per capita growth to CO2 emission from manufacturing industries and construction, arms import, commercial service export and coal rents, as well as a causal link between manufacturing industries, arms import, commercial service export and coal rent.

Likewise, the neoclassical view was endorsed in developing and developed countries, respectively the hypothesis impartiality. The main implication instigating from this research can be follow: 29 developed and developing countries should promote the use of renewable vitalities that are constantly restocked and which will not directly be diminished. Hence, the use of renewable vitalities will contribute to the decrease of GHGs emission.

Besides, 29 developed and developing countries may benefit from enhanced social stability, job opportunity by modernized technologies. Finally, as endeavors of future research, our aim to outspread the empirical analysis in order to verify and test the EKC hypothesis employing the environmental performance and encourage to developed countries to secure the environment especially for arms and huge manufacturing industries.

## Acknowledgment

The authors admiringly acknowledge the financial support of the National Natural Science Foundation of China No: 71371087 of Jiangsu University, as well as thanks to my supervisor Dr. YuSheng Kong about review and editing. Hina (Rawaa) is thankful to the excellence idea also appreciate for the study support in the Republic of China (Zhenjiang). I would like to thanks of my parents and co-author whose support me in different conceptual ideas and techniques. Finally, I have also benefited from generous support of my beloved Allah who give me competence and enthusiasm well to improve my skills.

## Footnotes

1 https://en.wikipedia.org/wiki/Neoclassical_economics

2 Retrieved 2016, from http://www.bp.com/content/dam/bp/pdf/energy-economics/statistical-review2016/bpstatistical-review-of-world-energy-2016-full-report.pdf.

3 *Sustainable Energy for all (SE4ALL)* 2013; Available from: https://data.worldbank.org.cn/indicator/EG.EGY.PRIM.PP.KD?View=chart

4 Climate change three chart shows the CO2 emission from 1850 to 2011. https://www.cgdev.org/media/who-caused-climate-change-historically

5 These 6 Countries Are Responsible For 60% Of CO2 Emissions [Press release]. Retrieved from https://www.businessinsider.com/these-6-countries-are-responsible-for-60-of-co2-emissions-2014-12

6 The China and USA deal on greenhouse gas emission growth by 2030, while its significant and also little effected on the global thermostat. The USA government estimates China doubling it emission by 2040 cause of major changes and reliant on fossil fuels for steel and electricity production. There was 2.6 billion tons CO2 emission in India with 1.2 billion population, 2 billion tons in Russia with 143.5 million population, 1.4 billion tons in Japan with 127 million population, 836 million tons in Germany with 80.6 million population in 2013

7 https://www.indexmundi.com/facts/indicators/EN.ATM.CO2E.SF.KT

8 *Carbon Dioxide Information Analysis Center*, in *Environmental Sciences Division*. CO2 emissions from solid fuel consumption (kt), USA Oak Ridge National Laboratory, Tennessee.

9 *Knoema*. 2014; Available from: https://knoema.com/atlas/topics/Environment/Emissions/CO2-emissions-from-gaseous-fuel-consumption?action=export&gadget=tranking-container.

10 Landfill, *Carbon Emissions from Waste Measured in EPA Greenhouse Gas Inventory*. 2010: USA. https://waste-management-world.com/a/carbon-emissions-from-waste-measured-in-epa-greenhouse-gas-inventory

11 According to Loesche, D, *The Carbon Age: 150 Years of CO2 Emissions*. 2018 https://www.statista.com/chart/13584/worldwide-carbon-emissions-from-fossil-fuel-consumption-and-cement-production/

12 The Carbon dioxide information analysis center (CDIAC), realized more than 400 billion metric tonnes in atmosphere from fossil consumption and especially production of cements since 1751. Also, the combustion of solid and liquid fossil fuel causes of 4^th^ of all CO2 which is 9.9 billion tones in 2014.

13 https://knoema.com/atlas/topics/Environment/Emissions/CO2-emissions-from-gaseous-fuel-consumption?action=export&gadget=tranking-container

